# *Iodidimonas*, a bacterium unable to degrade hydrocarbons, thrives in a bioreactor treating oil and gas produced water

**DOI:** 10.1101/2023.03.02.530844

**Authors:** Shwetha M. Acharya, Brandon C. Enalls, Peter J Walian, Brett D. Van Houghton, James S. Rosenblum, Tzahi Y. Cath, Susannah G. Tringe, Romy Chakraborty

## Abstract

*Iodidimonas* is a genus recently described in bioreactors treating oil and gas produced water and in iodide rich brines. Besides the ability to oxidize iodine, little is known about the metabolic capabilities that enable *Iodidimonas* sp. to occupy this unique ecological niche. We isolated, characterized, and sequenced three strains belonging to the *Iodidimonas* genus from the sludge of a membrane bioreactor treating produced water. We describe the genomic features of these isolates and compare them with the only other four isolate genomes reported from this genus, as well as a metagenome-assembled genome from the source bioreactor. To survive in the produced water, *Iodidimonas* isolates had several genes associated with mitigating salinity, heavy metal and organic compound stress. While the isolates could utilize a wide variety of carbon substrates, they failed to degrade aliphatic or aromatic hydrocarbons, consistent with the lack of genes associated with common hydrocarbon degradation pathways in their genomes. We hypothesize these microbes may lead a scavenging lifestyle in the bioreactor and similar iodide-rich brines.

**Importance:** Occupying a niche habitat and having few representative isolates, genus *Iodidimonas* is a relatively understudied Alphaproteobacterial group. This genus has garnered attention due to its ability to corrode pipes in iodine production facilities and generate iodinated organic compounds during treatment of oil and gas produced water. The iodinated organic compounds are likely to be carcinogenic and may pose issues with recycling the treated water. Hence, detailed characterization of the metabolic potential of these isolates is not only of economic importance, but also sheds light on adaptation of this microbe to its environmental niche.

## Introduction

Produced water (PW), the wastewater generated as a byproduct of oil and gas extraction, contains a wide variety of contaminants such as hydrocarbons and other organic compounds, salts, heavy metals and radionuclides (1–7). Biological pretreatment of PW has been shown to be effective in removal of organic contaminants (8–17). Specifically, hybrid physical and biological treatment systems like membrane bioreactors (MBRs) are gaining interest due to their modularity, low energy demand, and ability to retain biomass in the reactor (18) to catalyze removal of organic contaminants with minimal biofouling. However, limited knowledge about the microbes that form the basis for reactor functions can impede modeling and optimization of treatment processes (19).

Previously, we reported on the operation of an aerated MBR for a year treating Denver-Julesburg (DJ) Basin PW with a gradual increase in feed salinity from 30 to 100 g/L total dissolved solids (TDS) (20). Time-series monitoring of this reactor through 16S rRNA gene amplicon sequencing revealed that *Iodidimonas* was not only a dominant and persistent genus throughout the study duration but was also the first genus to colonize the reactor along with *Roseovarius* (20). A recent study of two membrane bioreactors treating shale gas wastewater in China also reported *Iodidimonas* as one of the dominant groups in the reactor (21).

*Iodidimonas* is a recently proposed genus in the class Alphaproteobacteria (GTDB Taxonomy: Bacteria; Proteobacteria; Alphaproteobacteria; Kordiimonadales; Iodidimonadaceae; *Iodidimonas*) and one of the few reported groups of bacteria capable of iodide oxidation (22). To date, only 9 isolates have been reported, out of which only 4 have publicly available genomes and very little is known about this genus (22–25). Members of this genus were first isolated from iodide-rich natural gas brines and iodide-amended surface seawater enrichments, at total iodine concentrations (1 mM) that were 2000-fold higher than that of typical seawater (0.5 µM) (23). Oxidation of iodide to molecular iodine (I_2_) and other volatile organic iodinated compounds is mediated by an extracellular multicopper oxidase named iodide-oxidizing enzyme (IOX) (26), originally discovered in *Iodidimonas* sp. Q-1. This IOX/iodide system is shown to have antimicrobial and sporicidal properties against wide variety of bacterial strains and spores (27). Because iodide-oxidation by an extracellular enzyme provides no metabolic energy to the microbe, it was proposed to be a strategy to kill other heterotrophic bacteria susceptible to iodine and gain dominance in iodide-rich waters (23, 28).

Growth of these iodide-oxidizing bacteria in iodide-rich brines can be undesirable. Members of the *Iodidimonas* genus were experimentally shown to be involved in corrosion of carbon steel pipes carrying brine to an iodine production facility (25). Enrichment of *Iodidimonas* and *Roseovarius* (another group of iodide-oxidizing bacteria) in biologically active filters treating oil and gas PW were implicated in the formation of toxic iodinated organic compounds in treated wastewater, which could negatively impact potential water recycling (29).

Members of this genus have been isolated from or detected mostly in subsurface-associated environments; many of these environments are reported to have high iodide concentrations and are also hydrocarbon-associated (natural gas brines/O&G PW) (20–25). While their ability to oxidize iodide to iodine has been investigated before, the mechanisms of survival in hydrocarbon associated environments have not been explored. Previously characterized isolates of *Iodidimonas* (i.e., *Iodidimonas muriae* C-3^T^, *Iodidimonas gelatinilytica sp*. Hi-2^T^, *Iodidimonas gelatinilytica sp*. Mie-1) have been described as strictly aerobic, moderately halophilic, chemoorganotrophic, and capable of iodide oxidation, but unable to grow on iodide as sole electron donor (22, 24).

In this study, we report on the isolation of three bacterial strains belonging to the *Iodidimonas* genus from the sludge of a lab-scale MBR treating PW. We demonstrated the phenotypic and genomic characteristics of our *Iodidimonas* isolates and compared their genomes to other members of this understudied genus as well as a composite metagenome-assembled genome (MAG) from the MBR. Based on our analysis of metabolic capabilities, we hypothesize the plausible reasons why these species successfully dominate the specific ecological niche of iodide-rich brine waters, including the MBR treating PW.

## RESULTS

### Phenotypic and biochemical characterization of three Iodidimonas isolates from an MBR

The three *Iodidimonas* strains were isolated on Marine Agar 2216 (MA) from sludge at time points when the concentration of total dissolved solids (TDS), a measure of salinity, of feed to the bioreactor was 27, 50 and 80 g/L (Fig 1). *Iodidimonas spp*. MBR-14, MBR-22, and MBR-55 have salinity tolerance ranges of 2-12%, 3-8% and 2-8% (w/v), respectively, and exhibit growth optima around 3% (Fig S1A). Typically, DJ Basin PW has TDS in the range of 20-40 g/L (equivalent to 2-4% w/v) but was supplemented with NaCl in the later stages of reactor operation to test the efficacy of the MBR at higher salinities (20). Strains MBR-22 and MBR-55 were isolated from the sludge when the feed was amended with NaCl but was still within the salinity tolerance range of the three strains. The three strains grew best in a pH range of 7.5-8.5 (Fig S1B) and had optimum growth in a temperature range of 20-30 °C (Fig S1C). Overall, strain MBR-14 had distinct growth pattern under salinity, temperature, and pH ranges compared to MBR-22 and MBR-55, which displayed similar growth curves.

**Fig 1.**
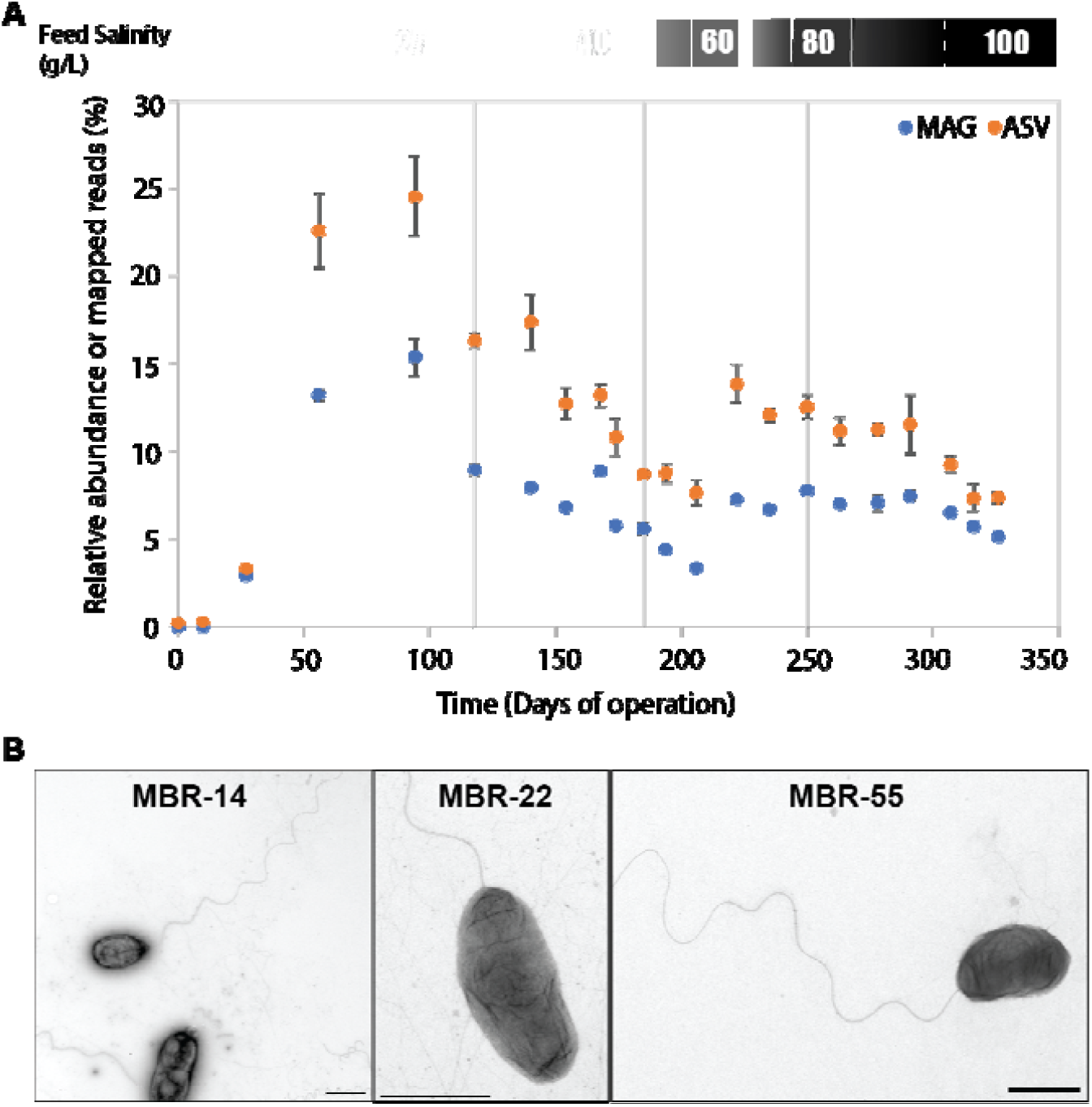
Plot showing (A) abundance of *Iodidimonas* MAG and ASV in the time-series data from MBR, (B) TEM micrographs of the three isolates. Scale bars represent 1 µm length in each TEM micrograph. Lines between the two panels link the isolates with the timepoints at which the were isolated.

The three strains grew on L-valine, L-glutamic acid, propionate, butyrate, glucose, maltose, sucrose, and thymidine (Table S2). They slightly differed in their C-substrate utilization and showed weak growth on L-alanine and citric acid. No growth or substrate utilization was observed when strains were inoculated in either aliphatic hydrocarbon (hexadecane) or an aromatic hydrocarbon mixture (mix of benzene, toluene, ethyl benzene, xylenes and naphthalene-BTEXN) (Fig S2). All the isolates exhibited capability for iodide oxidation as assessed by forming purple pigment around colonies on potassium iodide and starch-amended marine agar (MA) (Fig S3). The supernatants from the three strains displayed a slight emulsification effect on crude oil, but not on diesel or gasoline, indicating the absence of biosurfactant production (Fig S4). A thin and faint yellow color zone formed around the isolate colonies compared to the positive control in an overlaid-chromeazurol S (o-CAS) assay testing for production of siderophores, and hence siderophore production is unlikely (Fig S5).

These isolates exhibited darting motility when prepared as a wet mount and observed under phase-contrast microscope. Electron microscopy micrographs (Fig 1B) show that the three strains are rod-shaped with similar average sizes of 1.5-2 µm in length and 0.7-0.9 µm in width. Additionally, the three strains have a single flagellum (∼20 nm diameter) and multiple finer filaments (∼5-8 nm diameter), confirming the presence of appendages for bacterial motility. Phenotypic characteristics of newly isolated strains and previously reported *Iodidimonas* isolates are summarized in Table 1.

**Table 1.** Phenotypic features of isolates and members of genus *Iodidimonas*.

| Members* | MBR-14 | MBR-22 | MBR-55 | Q-1 | C-3<sup>T</sup> | Hi-2<sup>T</sup> | Mie-1 |
| --- | --- | --- | --- | --- | --- | --- | --- |
| Isolated from | PW-fed MBR (CO, USA) | PW-fed MBR (CO, USA) | PW-fed MBR (CO, USA) | Iodide-rich natural gas brine (Japan) | Iodide-rich natural gas brine (Japan) | Iodide-rich natural gas brine (Chiba, Japan) | Surface seawater (Mie, Japan) |
| Iodide-oxidation | + | + | + | + | + | + | + |
| Colony appearance | Circular, convex, translucent, entire margins, yellow, 0.5-1 mm dia, MA, 7 days | Circular, convex, translucent, entire margins, yellow, 0.5-1 mm dia, MA, 7 days | Circular, convex, translucent, entire margins, yellow, 0.5-1 mm dia, MA, 7 days | NA | Circular, convex, opaque, entire margins, creamy white, 0.5-1.5 mm dia, MA, 7 days | Circular, convex, opaque, entire margins, creamy white, 0.5-1.5 mm dia, MA, 7 days | Circular, convex, opaque, entire margins, creamy white, 0.5-1.5 mm dia, MA, 7 days |
| Cell size (µm) | 1.5 X 0.7 | 2.0 X 0.9 | 1.5 X 0.8 | NA | 1.3-3.6 | 0.3-0.5 X 1.2-4.4 | 0.3-0.5 X 1.2-4.4 |
| Hydrolysis of aesculin | NA | NA | NA | NA | + | - | - |
| Liquefaction of gelatin | NA | NA | NA | NA | - | + | + |
| pH range | 6.6-8.5 | 7.6-8.5 | 7-8.5 | NA | 4.5-8.5 | 5.3-9.0 | 4.5-8.5 |
| Temperature (°C) | 4-35 | 10-30 | 10-30 | NA | 4-40 | 4-40 | 4-40 |
| Salinity range (%) | 2-12 | 3-8 | 2-8 | NA | 0.5-9.0 | 0.5-9.0 | 0.5-9.0 |
| Other features | Aerobic, rods, darting motility | Aerobic, rods, darting motility | Aerobic, rods, darting motility | NA | Aerobic, rods, vibrating motility, no spore formation | Aerobic, rods, vibrating motility, no spore formation | Aerobic, rods, vibrating motility, no spore formation |
| Growth on C sources | L-valine, L-aspartic acid, L-glutamic acid, inosine, acetate, | L-valine, L-glutamic acid, propionate, glucose, | L-valine, L-glutamic acid, propionate, maltose, | NA | Yeast Extract (0.1% w/v), Soluble starch | Yeast Extract (0.1% w/v), L-arabinose and weakly | Yeast Extract (0.1% w/v), L-arabinose and weakly |
|  | propionate, glucose, maltose, sucrose, thymidine, butyrate, and weakly on citric acid, malic acid, L-isoleucine, L-leucine, L-alanine, pyruvate, succinate and L-histidine | maltose, sucrose, thymidine, butyrate, and weakly on citric acid, L-alanine, pyruvate, L-aspartic acid | sucrose, thymidine, butyrate, and weakly on glucose, citric acid, malic acid, L-aspartic acid, L-alanine, and L-histidine |  | (0.1%), weakly on acetate, D-glucose, maltose, sucrose (10 mM), capric acid in air | on D-mannose, D-mannitol, adipic acid and phenylacetic acid. | on D-mannose, D-mannitol, D-maltose, N-acetyl glucosamine and capric acid. |

\*Complete names: *Iodidimonas* sp. MBR-14, *Iodidimonas* sp. MBR-22, *Iodidimonas* sp. MBR-55, *Iodidimonas* sp. Q-1, *Iodidimonas muriae* C-3<sup>T</sup>, *Iodidimonas gelatinilytica* sp. Hi-2<sup>T</sup>, *Iodidimonas gelatinilytica* sp. Mie-1
NA- Not Available, MA- Marine Agar, PW- Produced water, MBR- Membrane Bioreactor

### Phylogenetic classification of the three isolates

The three isolates shared the same full-length 16S rRNA sequence, and therefore formed a single branch in a 16S rRNA gene tree (Fig 2a). The most closely related 16S gene is from *Iodidimonas sp*. Q-1 (formerly *Alpha proteobacterium* Q-1), in a small cluster containing multiple clone sequences from a geyser and an isolate from a natural gas brine. A separate *Iodidimonas* cluster included *Iodidimonas muriae* C-3, *I. gelatinilytica* Hi-2 and Mie-1, a seawater isolate, and clone sequences from iodide-rich brines and oyster shell (details of *Iodidimonas* sequences in Table S6). In a whole genome tree based on marker genes (Fig 2b), the three isolates also clustered with *Iodidimonas sp*. Q-1 and formed a monophyletic group with all other reported *Iodidimonas* genomes. The average nucleotide identity (ANI) among MBR strains was >99.9% while ANI between MBR strains and other *Iodidimonas* ranged from ∼78.5% for C-3, Hi-2, and Mie-1 to ∼95.8% for Q-1 (Table S3).

**Fig 2.**
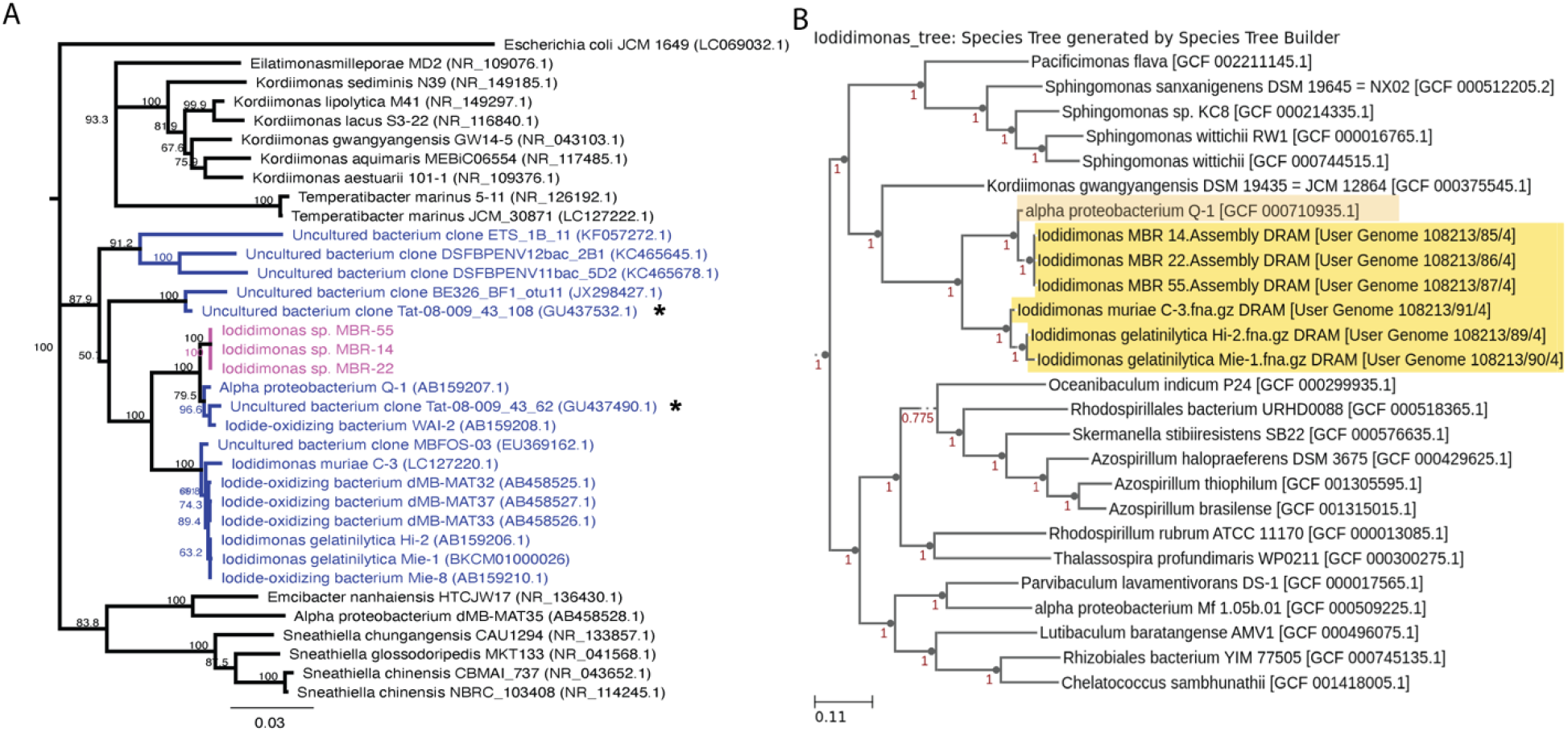
Phylogenetic trees based on (A) 16S rRNA sequences (Neighbor-Joining tree); isolates or clone sequences classified as genus *Iodidimonas* in SILVA database are marked in blue and the strains from this study in pink; (B) genome-based comparison (49 COGs) of the strains with closest sequenced relatives; *Iodidimonas* genomes are highlighted. 16S rRNA clone sequences marked with asterisk were selected as representative among other closely related clone sequences for easy visualization of this tree (A); full tree is shown in Fig S6.

### Abundance and genome-inferred growth rates of Iodidimonas in the reactor

Abundances of *Iodidimonas* (Fig 1A) in the MBR sludge during the experimental period was estimated using relative abundance in 16S rRNA amplicon data and % of raw metagenome reads mapped to the MAG and showed similar trends over time (Pearson’s correlation coefficient, *r* = 0.96). The index of replication (iRep), a proxy for replication rate based on genome coverage variation (30), was constant throughout the MBR operation period (Fig. S6). The average iRep value for all timepoints is about 1.13, indicating that approximately 13% of the *Iodidimonas* population is replicating at any given time. This suggests that *Iodidimonas* growth is largely indifferent to fluctuations of salinity and other environmental conditions within the bioreactor.

### Genome analysis of the three MBR isolates

Genome sizes of *Iodidimonas spp*. MBR-14, MBR-22, and MBR-55 varied slightly, with MBR-14 having the largest size of 3.04 Mbp (summarized in Table 2). All three isolate genomes had similar numbers of contigs and coding sequences (CDS), and G+C content. The MAG generated from MBR sludge metagenomes had, as anticipated, a smaller size of 2.78 Mbp in 23 contigs. While each of the three isolate genomes had a single 16S-23S-5S rRNA operon and 47 tRNAs, the MAG lacked the rRNA operon and had fewer tRNA genes.

**Table 2.** Summary of genome features of three MBR isolates and MAG.

| Genome features | MBR-14 | MBR-22 | MBR-55 | MAG |
| --- | --- | --- | --- | --- |
| Size (Mbp) | 3.04 | 2.97 | 2.98 | 2.78 |
| GC content | 56.6 | 56.43 | 56.43 | 56.3 |
| Number of contigs | 46 | 43 | 45 | 23 |
| Completeness (%) | 99.57 | 99.57 | 99.57 | 97.61 |
| Contamination (%) | 0 | 0 | 0 | 0 |
| No. of CDS | 2621 | 2557 | 2567 | 2370 |
| No. of rRNA operons<br>(16S-23S-5S) | 1 | 1 | 1 | 0 |
| No. of tRNAs | 47 | 47 | 47 | 41 |

All major genome features described below are tabulated in Supplementary Tables S4 and S5. For nutrient acquisition and adaptation, several types of transporters (ATP-binding cassettes (ABC)-type, Tripartite ATP-independent periplasmic (TRAP)-type, antiporter, Major facilitator superfamily proteins (MFS) and symporters) components were predicted. Type II Secretion system (T2SS) genes for secretion of extracellular proteins were present, which are critical for adaptation to the environment and biopolymer recycling (31). Several carbohydrate active enzymes (CAZy) and peptidases were found, which play a critical role in carbon metabolism. No siderophore genes were predicted in the genomes, but a TonB-ExbB-ExbD system of siderophore uptake was present. This agrees with the absence of siderophore production as indicated by o-CAS assay, suggesting the presence of alternate mechanisms of iron acquisition. Genes for hemin uptake (*hmuPSTV*) were predicted in all three MBR genomes. No major surfactant genes were found in the genomes, in congruence with the results of an emulsification activity test. A gene cluster associated with the iodide oxidizing enzyme IOX (*ioxABCDEF*), previously reported from *Iodidimonas* sp. Q-1, was also present in these strains. Gene *ioxA* was annotated as a multicopper oxidase and *ioxB, ioxD*, and *ioxF* were homologous to the SCO1/SenC family of proteins, and genes belonging to general secretion pathways were adjacent to the *iox* gene cluster, as reported in other *Iodidimonas* strains (26, 32).

All the isolates had complete or near-complete pathways predicted for glycolysis, tricarboxylic acid (TCA) cycle, pentose-phosphate pathway, and Entner-Doudoroff pathway. No genes associated with known hydrocarbon degradation pathways were found in any *Iodidimonas* genomes, in agreement with the lack of hydrocarbon degradation in our experiments. Various proteins involved in aerobic oxidative phosphorylation (cytochrome C oxidase) were predicted; however, there is no genomic evidence indicating utilization of electron acceptors other than oxygen.

Production and transport of organic osmolytes along with regulation of inorganic ions confer salinity tolerance in prokaryotes. Among organic osmolytes, the *Iodidimonas* strains harbor genes for synthesis and/or transport of betaine, glutamine, glutamate, and proline, but not ectoine. Homologs of genes for betaine transport (*betT*) and biosynthesis of proline (*proABC*), glutamate (*gltB*) and glutamine (*glnA*) were predicted in all three genomes. Potassium transport systems (*trkA, trkH*), a chloride ion channel, and several Na^+^/Ca^2+^/K^+^-H^+^ antiporter associated genes predicted could be involved in the regulation of ionic transport across the cell membrane.

Besides sodium and chloride ions, major ions found in PW include boron, barium, calcium, iron, potassium, lithium, magnesium, silicon, strontium, fluoride, bromide, and sulfate; most of these passed through the MBR unaltered in concentration (20). Phosphate and nitrate were below detection limits throughout the study. Other ions detected at lower concentrations (<1ppm) include zinc, manganese, copper, and arsenic. Iodide concentration in the bioreactors was not measured in this study but iodide have been detected in the range 40-53 mg/L (0.32-0.42 mM) in DJ Basin PW (29). Several genes associated with transport, accumulation, and regulation of copper, zinc and iron were predicted; there were a few genes associated with arsenic regulation, transport and transformation. Homologs of genes associated with fluoride exporter (*crcB*), responsible for reduction of fluoride concentrations inside the bacterial cells (33), were also detected in these strains. The presence of heavy metals is known to trigger oxidative stress responses in bacteria (34, 35). Genes to mitigate oxidative stress and damage include thioredoxin, glutaredoxin, superoxide dismutase and peroxidase were present in all the three strains along with several heat shock, cold shock, and phage shock proteins and DNA repair mechanisms. Phage shock proteins help faster recovery of cells from different agents that impact the cell membrane (36). All three genomes harbored a homolog for Cyclopropane-fatty-acyl-phospholipid synthase, which produces cyclopropane-fatty-acid, a molecule that is known to provide protection against desiccation (37). Additionally, ABC-type transporters that are involved in resistance to organic solvents (*mlaCDE*) were also predicted along with a few other genes (*ttgE, tolC*) known to confer resistance to organic compounds.

Near-complete gene sets were identified for flagella and pilus assembly alongwith chemotaxis proteins, which are critical for motility as well as adhesion. Presence of pili and flagella was also verified in electron microscopy images (Fig 1B). A few genes associated with exopolysaccharide/ biofilm formation were present, even though *eps* genes were not predicted. While we did not specifically perform biofilm formation assays on the three strains, they did exhibit clumping and formed biofilm-like structures after a few days of growth in all the different growth media.

Other interesting genomic features that may not necessarily relate to the survival of these three strains in the reactor. Photosynthetic gene clusters (photosystem II) are present, but pathways for autotrophic CO_2_ fixation are missing, suggesting that these strains may be aerobic anoxygenic phototrophs. Phage signatures were predicted as well as toxin-antitoxin and CRISPR-Cas defense systems. Predicted biosynthetic gene clusters (BGCs) include a carotenoid, terpene, homoserine lactones, RiPP-like, and N-acetylglutaminylglutamine amide (NAGGN). NAGGN has been shown to be an important osmolyte, protecting bacterial cells against desiccation and salinity (38).

### Comparative genomics with other members of the Iodidimonas genus

The four previously reported *Iodidimonas* genomes have an average size of 2.94 Mb and G+C content of 55.6% (32), comparable to the isolates described in our study. Comparative genomics was performed based on presence or absence of gene clusters and homogeneity indices of the aligned genes (Fig 3). Homogeneity indices calculated by anvi’o are a combination of geometric homogeneity (gap/residue patterns within the gene cluster) and functional homogeneity (conservation of aligned residues across genes) of the gene clusters (https://merenlab.org/2016/11/08/pangenomics-v2/#inferring-the-homogeneity-of-gene-clusters). There were 1,629 and 928 genes that were identified as core (present in all genomes) and singleton (present in only one genome) genes, respectively. Accessory genes were split into C3-cluster specific (present only in strains C-3, Hi-2, and Mie-1), MBR-Q1 cluster specific (present in all strains from MBR and Q-1), MBR-specific (found only in MBR strains) and other accessory genes (present in more than one genome with no specific pattern) which totaled to 574, 559, 194, and 472 genes, respectively.

**Fig 3.**
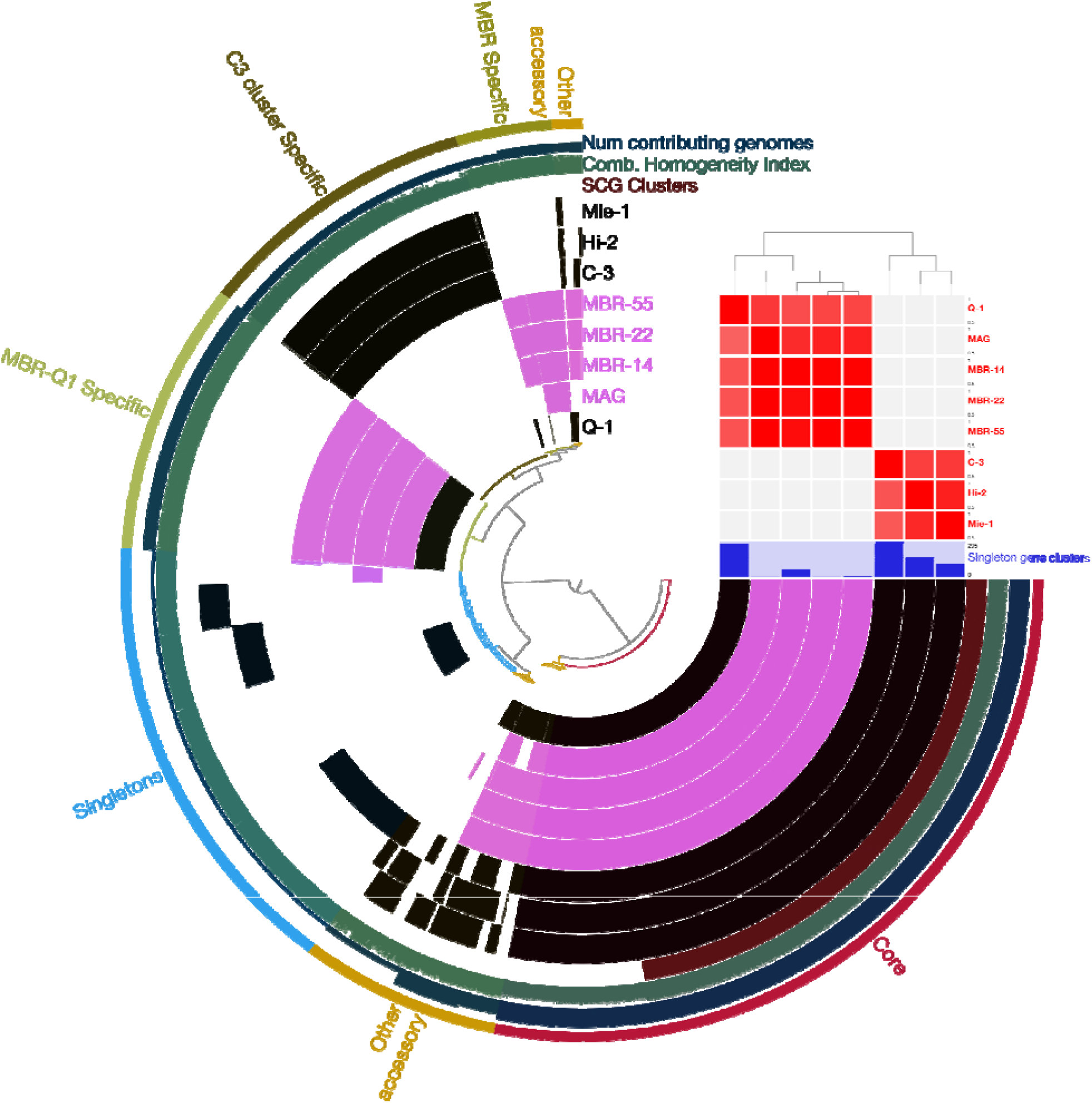
Circular plots comparing genomes from the *Iodidimonas* genus based on gene cluster presence/absence (Euclidean distance and ward linkage). The isolates and MAG presented in this study are shown in pink. The core (present in all genomes), accessory (present in two or more genomes) and singleton (present in one genome) gene clusters are labeled outside the outer ring. Accessory genes are further binned into C3-cluster specific, MBR-specific, MBR-Q1-cluster specific, and other accessory genes. The matrix on the top right shows ANI distances between various genomes with darker color indicating higher similarity.

A large proportion of core genes belong to single-copy gene clusters that are conserved across the bacterial domain. The MBR-Q1 cluster specific genes include the entire photosynthetic gene cluster; one gene cluster of Type II secretion systems; a few genes associated with flagellar and pilus assembly; several ABC-transporters and extracellular binding proteins; beta-lactamase; polysaccharide synthesis; and other transport proteins. MBR-specific accessory genes included one of the phage signatures (set-2) and some CRISPR-associated elements; toxin-antitoxin components; the entire urease cluster (*<u>ureABCDEF</u>*); several DNA binding and repair proteins; a few genes associated with flagellar assembly; antibiotic resistance proteins; heavy-metal/metal binding domains; subtilase; and other structural and regulatory proteins. The differences in flagellar and chemotaxis proteins may explain why *Iodidimonas* spp. MBR-14, MBR-22, and MBR-55 are motile with visible flagella while *Iodidimonas muriae* C-3, *I. gelatinilytica* Hi-2, and *I. gelatinilytica* Mie-1 are non-motile based on previous reports. C3-cluster specific accessory genes contain several ABC-transporters and extracellular binding proteins, beta-lactamase, polysaccharide synthesis, and transport proteins; many of these share common functions but not sequence homology with MBR-Q1 cluster genes.

The *Iodidimonas* MAG generated from metagenomes captured a large portion of the genes conserved among the MBR isolates. Among the three isolates from the reactor, strain MBR-14 has the highest number of singletons, including 63 genes located on a single contig (contig #15). The singleton genes include genes associated with copper/metal binding, copper transport and resistance genes, regulatory proteins and another multicopper oxidase gene. These MBR-14 specific genes were flanked by a gene cluster containing conjugal transfer proteins (*trbCDEFGJL*) and may indicate the acquisition of this genomic material through horizontal gene transfer. These conjugal transfer proteins are not found in the strains MBR-22 and MBR-55 but are shared with strain Q-1. Strains MBR-55 and MBR-22 had 9 and 1 singleton genes, respectively; however, most of these genes were unannotated. One interesting singleton from strain MBR-55 was predicted to be gene *qacE* whose product is involved in transport of cations and cationic drugs. This gene is implicated in resistance to quaternary ammonium compounds (QACs), phenanthridines and xanthenes, some of which are reported to be found in PW (39).

## DISCUSSION

### Iodidimonas is abundant, persistent, and yet slow growing in the bioreactor

*Iodidimonas* and *Roseovarius* were the first dominant groups to colonize the bioreactor as PW was introduced and the salinity of the feed increased during the acclimation phase. This was evident when the relative abundance of an ASV matching the *Iodidimonas* isolates slowly increased from 0 to 4 to almost 25% when the feed salinity increased from 1.5 g/L to 20 g/L to 27 g/L. This conformed with the growth of MBR strains in the lab only when the salinity was above 2% (w/v), with an optimum around 3% (w/v). The *Iodidimonas* ASV and MAG from the MBR follow the same temporal abundance trends, thus providing greater confidence that the change in abundance is due to an actual change in community composition.

Surprisingly, the replication index for the MAG stayed constant at 1.13 throughout the reactor run despite large changes in feed salinity. A similar uniform distribution of replication rates for activated sludge MAGs during periods of disturbance and stable operation was observed in another study (40), and the authors inferred that growth rate was not a primary driver for bacterial selection under disturbance in that system. Further, the design of the MBR, an aerated tank where the effluent is removed via an ultrafiltration membrane, results in retention of all the biomass - whether introduced through feed, actively growing, quiescent, or dead - in the reactor. However, the cultivability of *Iodidimonas* at different timepoints during the reactor run implies that at least a fraction of the *Iodidimonas* population was alive and persisted in the reactor and was not merely detected due to accumulation of dead or nonreplicating cells.

### Connecting genotype and phenotype to survival in the reactor

pH (7.5-8.5) and temperature optima (20-30 °C) for all three strains were congruent with the conditions at which the reactor was operated (pH 7.5-8.5, 20 °C), making the MBR a conducive environment for proliferation of these organisms. MBR-14 has slightly different pH, salinity, and temperature growth curves and carbon substrate utilization pattern compared to MBR-22 and MBR-55, possibly due to a number of singleton genes in MBR-14. Another key reactor parameter that is often reported to promote growth of *Iodidimonas* is aeration (21, 29, 32). The influence of aeration could be two-fold, by a) enhancing aerobic metabolism (oxidative phosphorylation machinery) thus eliminating anaerobes present in feed, and b) killing of iodine-sensitive, competing bacteria through secretion of iodide oxidizing enzyme, which in turn oxidizes iodide only in the presence of oxygen. Another key factor is the presence of iodide in the PW, which is reported in the range of 40-53 mg/L in DJ Basin PW (29).

PW is not just an iodide-rich brine; it contains several other contaminants like heavy metals and inorganic and organic compounds which originate from the geological formation, as well as chemicals added during the drilling and operation of the oil and gas wells. The presence of a large number of putative heavy metal, salinity (osmoprotection), and organic solvent tolerance genes along with genes for mitigating oxidative stress may explain survival of *Iodidimonas* in the reactor. In addition, these genes may confer the ability to counteract the negative effects of iodine generated via iodide oxidation.

Genome analysis revealed that the *Iodidimonas* strains had several transporters, CAZy enzymes, and peptidases for acquisition and degradation of nutrients from the environment. Experiments confirmed that the strains could utilize a few simple sugars, amino acids, organic acids, and nucleotides as sole sources of carbon and energy. However, biochemical and genomic characterization revealed a lack of degradation pathways for commonly reported hydrocarbons, which is surprising given that this genus is abundant in hydrocarbon-associated environments. We hypothesize that the *Iodidimonas* population in the reactor obtains energy through one or more of the following methods: a) the killing of iodine-susceptible bacteria by the secreted IOX and feeding on their lysed necromass, b) cross-feeding on metabolites/substrates released by other sludge bacteria, and/or c) presence of novel pathways for degradation of hydrocarbons (lesser-known hydrocarbons present in the reactor with not so well-characterized degradation pathways). In a previous study, genome-resolved metagenomics identified that *Saccharibacteria*, a highly abundant genus in hydrocarbon-amended enrichments, lacks hydrocarbon degradation pathways and leads a scavenging lifestyle dependent on hydrocarbon-degrading hosts (41). Similarly, *Iodidimonas* may also act as a carbon sink by scavenging from the environment instead of actively degrading hydrocarbons, though *Iodidimonas* does not exhibit amino acid and nucleotide auxotrophies like the *Saccharibacterial* MAGs and is capable of survival without a host.

### Comparative genomics yields new insight about the genus Iodidimonas

The ANI between the MBR strains is greater than or equal to 99.98%, which implies that these strains can be categorized into a single species and perhaps even a common subtype. Phylogenetic analysis revealed that while all the *Iodidimonas* genomes formed a monophyletic cluster, there were distinct genetic groups within the genus - MBR isolates clustered closely with *Iodidimonas* sp. Q-1 while *I. muriae* C-3, *I. gelatinilytica* Hi-2, and *I. gelatinilytica* Mie-1 formed a separate cluster. This was also observed with ANI and alignment fraction (AF) values for these genomes. Comparative genomics based on presence/absence of gene clusters further revealed that the MBR isolates shared many genomic features including the massive photosynthetic gene cluster with *Iodidimonas sp*. Q-1 (42) while C3-cluster also had a fair share of common genes unique to the cluster. One set of phage signatures was unique to MBR isolates, indicating the influence of mobile genetic elements present in the local environment in shaping the genome. Despite this, more than 60% of the genes were conserved across the genus, constituting the core genes, which are perhaps critical for the survival of these members in iodide-rich brines.

Through this study, we have elucidated survival strategies of the understudied Alphaproteobacterial genus *Iodidimonas* in hydrocarbon-rich environments. Genome and biochemical analyses of the three *Iodidimonas* strains we isolated from a bioreactor treating produced water uncovered various strategies used by these microbes to survive in the produced water. Surprisingly, these isolates lacked hydrocarbon degradation pathways and failed to degrade hydrocarbons in lab assays, despite belonging to a highly abundant and persistent genus in the reactor. Hence, *Iodidimonas* may utilize necromass of iodine-sensitive bacteria and/or obtain substrates/metabolites from other bacterial groups in the reactor rather than through hydrocarbon degradation.

## MATERIAL AND METHODS

### Bacterial isolation and Whole Genome Sequencing

Sludge samples from a membrane bioreactor treating produced water at the Colorado School of Mines, Golden CO (20) were periodically collected in sterile, 50 mL centrifuge tubes and shipped overnight to LBNL on ice. Sludge samples were thoroughly mixed, serially diluted and spread on Marine Agar 2216 plates (Difco, BD Biosciences, USA) and incubated at 30 °C upon receipt. Colonies were picked and purified through further streaking, grown to log-phase in Marine broth (MB), purity and identity of isolates were determined by 16S rRNA sequencing, and cultures were preserved in 25% glycerol stock for long-term storage. The strains described in this study *-Iodidimonas sp*. MBR-14, *Iodidimonas sp*. MBR-22, and *Iodidimonas sp*. MBR-55 - were isolated on MA from sludge collected on Day 118, Day 185, and Day 250 of the reactor operation when TDS of feed was 27, 50, and 80 g/L, respectively.

DNA extraction and species identification was performed as described previously (43). Briefly, genomic DNA was extracted from cell pellets using PureLink Genomic DNA Mini kit (Thermo Fisher), 16S rRNA gene was amplified with universal bacterial primers (27F and 1492R) (44) and Sanger sequencing was performed. Forward and reverse reads were merged using Geneious Prime 2020.2.5 (https://www.geneious.com) to generate consensus sequence and queried against SILVA database (45, 46) for taxonomic classification.

Whole genome sequencing was performed at Novogene Corp. using Illumina NovaSeq 6000 platform (PE150). Raw reads were assessed for quality using FastQC v.0.11.9 (47), adaptor-trimmed and quality filtered through Trimmomatic v.0.36 (48) and assembled using default parameters with SPAdes v.3.15.3 (49) on KBase (50). Quality of assembled genomes was checked with Quast v. 4.4 (51) and CheckM v. 1.0.18 (52), also implemented in KBase.

### Biochemical characterization-pH, salinity, temperature growth curves

The bacterial strains were grown to mid-log phase in MB, washed thrice and resuspended in PBS buffer to be used as inoculum. For salinity tests, R2A broth was amended with NaCl to generate the following concentrations: 0, 0.5, 1, 2, 3, 4, 6, 8, 10, 12, 16, and 20% (w/v) NaCl. pH tests were conducted on R2A media amended with 3% (w/v) NaCl and appropriate buffers to create pH levels in the range of 3.4-9.16. Both salinity and pH tests were carried out in 48-well plates with 10% inoculum, incubated in dark at 30 °C with 200 rpm shaking. Optical density at 600 nm (OD_600_) was measured at regular intervals. Temperature growth curves were determined on R2A media with 3% NaCl, incubated at different temperatures of 4, 10, 20, 25, 30, 35, and 40 °C. 10% inoculum was added to culture tubes with media and was sampled at regular intervals for OD_600_ measurement. Growth rates were calculated using natural logarithm of OD_600_ versus time during log-phase.

### C-substrate utilization, auxiliary activities, and electron microscopy

Growth of the isolates on 50 different carbon substrates (Table S1) in artificial pore water media (APM; recipe in Supplementary material) was tested. 96 well plates containing APM with individual carbon substrates at 10 mM concentration and 10% (v/v) inoculum (triplicate plates for each strain) were incubated in dark, 30 °C, 120 rpm for 5 days. OD_600_ was measured daily and OD_600_ at the end of 5 days was used to determine growth on the substrates by comparing against uninoculated, negative controls.

Hydrocarbon degradation potential was assessed by monitoring utilization of BTEXN or hexadecane in APM coupled with cell enumeration using flow cytometry; auxiliary activities tested include ability to oxidize iodine, emulsify hydrocarbons and produce siderophores for iron acquisition (23, 53, 54), methods for these tests are described in Supplementary material alongwith that of electron microscopy imaging.

### Iodidimonas ASV, metagenome assembled genome (MAG) and their abundance and growth rates in the reactor

Experimental design and sampling of the MBR sludge at 22 different timepoints, and subsequent 16S amplicon sequencing, are described in detail in (20). Briefly, sludge samples from each timepoint were split into three 1.5 mL replicates and extracted using Qiagen DNeasy PowerLyzer PowerSoil kit as per manufacturer’s instructions. 16S rRNA gene amplicon sequencing was carried out at Novogene using universal bacterial primers 515F/806R (55) and processed via QIIME2 (56) and taxonomically classified against SILVA database v. 138 (45, 57). An Amplicon Sequence Variant (ASV) with 100% identity to the isolate full-length 16S sequences was identified through BLAST+ (58) and relative abundance of this ASV was determined in all samples.

Shotgun sequencing, assembly and binning of the metagenomes was carried out at the Joint Genome Institute (59). Raw reads were filtered for any artifacts or contaminants using BBTools and corrected using bbcms v.38.86 (sourceforge.net/projects/bbmap/). This readset was assembled using metaSPAdes v. 3.14.1 (60) and binning was performed using MetaBAT v2:2.15 (61) and quality of the bins was determined with CheckM v1.1.3 (52).Medium and high-quality MAGs from the sludge metagenomes at all timepoints were further selected and dereplicated using dRep (62). Among the representative genomes generated by dRep, a MAG associated with the GTDB-tk classification ‘*Iodidimonas* sp000710935’ was selected for further analysis (63). Percentage of raw reads mapped to the MAG at various timepoints using BBMap (sourceforge.net/projects/bbmap/) was used as a proxy for abundance of this MAG. For calculation of bacterial replication rates from metagenomes, raw reads from each metagenome were mapped to the MAG and sorted using bowtie2 (64) to generate sam files; these sorted sam files were used to calculate index of replication using iRep (30) based on coverage trend along the MAG.

### Genome analysis and Comparative genomics

Phylogenetic trees based on 16S rRNA sequences and whole genome sequences were generated to infer evolutionary relationships among the studied strains and their closest neighbors. Full length or near-full length 16S rRNA sequences were chosen based on previous studies (22, 24) and sequences (>1200 bp length) classified as belonging to *Iodidimonas* genus in the SILVA database (45, 46). All the selected 16S rRNA gene sequences were aligned using MUSCLE (65) and trees were generated using Neighbor-Joining algorithm (66) in Geneious Prime 2020.2.5 (https://www.geneious.com), 1000 bootstraps, using E. coli as an outgroup. For the whole genome tree, DRAM annotated genomes of MAG and the three study isolates were inserted into a phylogenetic tree with 20 closest RefSeq (67) genomes using “Insert Set of Genomes Into Species Tree” tool in KBase based on 49 single copy clusters of orthologous groups (COGs).

Assembled genomes of three strains from this study, the *Iodidimonas* MAG and four publicly available genomes of *Iodidimonas* were annotated using Eggnog-mapper v.2.1.3 (68). All the annotated genomes were then imported into anvi’o (69) and pangenome analysis was performed with parameters --minbit 0.5 --mcl-inflation 10. The circular plot generated by anvi’o was manually binned as core, accessory, and singleton gene clusters. Accessory gene clusters were further broken down into MBR-specific, MBR-Q1 cluster specific, C3-cluster specific and other accessory gene clusters to identify differences between the *Iodidimonas* groups. Average nucleotide identity across genomes was calculated with tool ‘PyANI’ in anvi’o and displayed as a matrix on the top right corner of the plot.

Additionally, all *Iodidimonas* genomes were compared against each other in a pairwise manner using FastANI (70) implemented in KBase. Genome alignment fraction (AF) was calculated by dividing the count of bidirectional fragment mappings by the number of total query fragment. Average Nucleotide Index (ANI) and AF were averaged for reciprocal comparisons. Predicted amino acid sequences from the three study isolates were further queried against additional, curated databases: BacMet (71) for heavy metal resistance genes, BioSurfDB (72) for surfactant genes and antiSMASH (73) using default parameters for detection of biosynthetic gene clusters (BGC).

## Acknowledgements

This study was supported by the Laboratory Directed Research and Development (LDRD) funding from Berkeley Lab under award No. 20-087. The work (DOI: 10.46936/10.25585/60001319) conducted by the U.S. Department of Energy Joint Genome Institute (https://ror.org/04xm1d337), a DOE Office of Science User Facility, is supported by the Office of Science of the U.S. Department of Energy operated under Contract No. DE-AC02-05CH11231 and used resources of the National Energy Research Scientific Computing Center, which is supported by the Office of Science of the US DOE (contract no. DE-AC02– 05CH11231).

Authors would like to thank Omolara T. Aladesanmi, Xuanqi Chen, Eunice Tsang, Shruthi Reddy and Rachael Peng for laboratory assistance. The authors also thank the ZOMA Foundation for support of the research at the Colorado School of Mines.

## References

1. Cluff MA, Hartsock A, MacRae JD, Carter K, Mouser PJ. 2014. Temporal changes in microbial ecology and geochemistry in produced water from hydraulically fractured Marcellus shale gas wells. Environ Sci Technol 48:6508–6517.

2. Orem W, Tatu C, Varonka M, Lerch H, Bates A, Engle M, Crosby L, McIntosh J. 2014. Organic substances in produced and formation water from unconventional natural gas extraction in coal and shale. International Journal of Coal Geology 126:20–31.

3. Khan NA, Engle M, Dungan B, Holguin FO, Xu P, Carroll KC. 2016. Volatile-organic molecular characterization of shale-oil produced water from the Permian Basin. Chemosphere 148:126–136.

4. Camarillo MK, Domen JK, Stringfellow WT. 2016. Physical-chemical evaluation of hydraulic fracturing chemicals in the context of produced water treatment. J Environ Manage 183:164–174.

5. Kondash AJ, Albright E, Vengosh A. 2017. Quantity of flowback and produced waters from unconventional oil and gas exploration. Sci Total Environ 574:314–321.

6. Rosenblum J, Thurman EM, Ferrer I, Aiken G, Linden KG. 2017. Organic Chemical Characterization and Mass Balance of a Hydraulically Fractured Well: From Fracturing Fluid to Produced Water over 405 Days. Environ Sci Technol 51:14006–14015.

7. Rosenblum J, Nelson AW, Ruyle B, Schultz MK, Ryan JN, Linden KG. 2017. Temporal characterization of flowback and produced water quality from a hydraulically fractured oil and gas well. Sci Total Environ 596–597:369–377.

8. Wang Z, Pan F, Hesham AE-L, Gao Y, Zhang Y, Yang M. 2015. Impacts of produced water origin on bacterial community structures of activated sludge. J Environ Sci (China) 37:192–199.

9. Pendashteh AR, Fakhru’l-Razi A, Chuah TG, Radiah ABD, Madaeni SS, Zurina ZA. 2010. Biological treatment of produced water in a sequencing batch reactor by a consortium of isolated halophilic microorganisms. Environ Technol 31:1229–1239.

10. Zhang X, Chen A, Zhang D, Kou S, Lu P. 2018. The treatment of flowback water in a sequencing batch reactor with aerobic granular sludge: Performance and microbial community structure. Chemosphere 211:1065–1072.

11. Sharghi EA, Bonakdarpour B, Roustazade P, Amoozegar MA, Rabbani AR. 2013. The biological treatment of high salinity synthetic oilfield produced water in a submerged membrane bioreactor using a halophilic bacterial consortium. J Chem Technol Biotechnol https://doi.org/10.1002/jctb.4061.

12. Kose Mutlu B, Ozgun H, Ersahin ME, Kaya R, Eliduzgun S, Altinbas M, Kinaci C, Koyuncu I. 2019. Impact of salinity on the population dynamics of microorganisms in a membrane bioreactor treating produced water. Sci Total Environ 646:1080–1089.

13. Riley SM, Ahoor DC, Cath TY. 2018. Enhanced biofiltration of O&G produced water comparing granular activated carbon and nutrients. Sci Total Environ 640–641:419–428.

14. Riley SM, Oliveira JMS, Regnery J, Cath TY. 2016. Hybrid membrane bio-systems for sustainable treatment of oil and gas produced water and fracturing flowback water. Separation and Purification Technology 171:297–311.

15. Freedman DE, Riley SM, Jones ZL, Rosenblum JS, Sharp JO, Spear JR, Cath TY. 2017. Biologically active filtration for fracturing flowback and produced water treatment. Journal of Water Process Engineering 18:29–40.

16. Shrestha N, Chilkoor G, Wilder J, Ren ZJ, Gadhamshetty V. 2018. Comparative performances of microbial capacitive deionization cell and microbial fuel cell fed with produced water from the Bakken shale. Bioelectrochemistry 121:56–64.

17. Camarillo MK, Stringfellow WT. 2018. Biological treatment of oil and gas produced water: a review and meta-analysis. Clean Techn Environ Policy 20:1127–1146.

18. Chang H, Li T, Liu B, Vidic RD, Elimelech M, Crittenden JC. 2019. Potential and implemented membrane-based technologies for the treatment and reuse of flowback and produced water from shale gas and oil plays: A review. Desalination 455:34–57.

19. Acharya SM, Chakraborty R, Tringe SG. 2020. Emerging trends in biological treatment of wastewater from unconventional oil and gas extraction. Front Microbiol 11:569019.

20. Van Houghton BD, Acharya SM, Rosenblum JS, Chakraborty R, Tringe SG, Cath TY. 2022. Membrane Bioreactor Pretreatment of High-Salinity O&G Produced Water. ACS EST Water 2:484–494.

21. Liu X, Tang P, Liu Y, Xie W, Chen C, Li T, He Q, Bao J, Tiraferri A, Liu B. 2022. Efficient removal of organic compounds from shale gas wastewater by coupled ozonation and moving-bed-biofilm submerged membrane bioreactor. Bioresour Technol 344:126191.

22. Iino T, Ohkuma M, Kamagata Y, Amachi S. 2016. Iodidimonas muriae gen. nov., sp. nov., an aerobic iodide-oxidizing bacterium isolated from brine of a natural gas and iodine recovery facility, and proposals of Iodidimonadaceae fam. nov., Iodidimonadales ord. nov., Emcibacteraceae fam. nov. and Emcibacterales ord. nov. Int J Syst Evol Microbiol 66:5016–5022.

23. Amachi S, Muramatsu Y, Akiyama Y, Miyazaki K, Yoshiki S, Hanada S, Kamagata Y, Ban-nai T, Shinoyama H, Fujii T. 2005. Isolation of iodide-oxidizing bacteria from iodide-rich natural gas brines and seawaters. Microb Ecol 49:547–557.

24. Iino T, Oshima K, Hattori M, Ohkuma M, Amachi S. 2021. Iodidimonas gelatinilytica sp. nov., aerobic iodide-oxidizing bacteria isolated from brine water and surface seawater. Antonie Van Leeuwenhoek 114:625–631.

25. Wakai S, Ito K, Iino T, Tomoe Y, Mori K, Harayama S. 2014. Corrosion of iron by iodide-oxidizing bacteria isolated from brine in an iodine production facility. Microb Ecol 68:519–527.

26. Suzuki M, Eda Y, Ohsawa S, Kanesaki Y, Yoshikawa H, Tanaka K, Muramatsu Y, Yoshikawa J, Sato I, Fujii T, Amachi S. 2012. Iodide oxidation by a novel multicopper oxidase from the alphaproteobacterium strain Q-1. Appl Environ Microbiol 78:3941–3949.

27. Yuliana T, Ebihara K, Suzuki M, Shimonaka C, Amachi S. 2015. A novel enzyme-based antimicrobial system comprising iodide and a multicopper oxidase isolated from Alphaproteobacterium strain Q-1. Appl Microbiol Biotechnol 99:10011–10018.

28. Arakawa Y, Akiyama Y, Furukawa H, Suda W, Amachi S. 2012. Growth stimulation of iodide-oxidizing α-Proteobacteria in iodide-rich environments. Microb Ecol 63:522–531.

29. Almaraz N, Regnery J, Vanzin GF, Riley SM, Ahoor DC, Cath TY. 2020. Emergence and fate of volatile iodinated organic compounds during biological treatment of oil and gas produced water. Sci Total Environ 699:134202.

30. Brown CT, Olm MR, Thomas BC, Banfield JF. 2016. Measurement of bacterial replication rates in microbial communities. Nat Biotechnol 34:1256–1263.

31. Nivaskumar M, Francetic O. 2014. Type II secretion system: a magic beanstalk or a protein escalator. Biochim Biophys Acta 1843:1568–1577.

32. Amachi S, Iino T. 2022. The genus iodidimonas: from its discovery to potential applications. Microorganisms 10:1661.

33. Calero P, Gurdo N, Nikel PI. 2022. Role of the CrcB transporter of Pseudomonas putida in the multi-level stress response elicited by mineral fluoride. Environ Microbiol https://doi.org/10.1111/1462-2920.16110.

34. Chasteen TG, Fuentes DE, Tantaleán JC, Vásquez CC. 2009. Tellurite: history, oxidative stress, and molecular mechanisms of resistance. FEMS Microbiol Rev 33:820–832.

35. Pal A, Bhattacharjee S, Saha J, Sarkar M, Mandal P. 2022. Bacterial survival strategies and responses under heavy metal stress: a comprehensive overview. Crit Rev Microbiol 48:327–355.

36. Joly N, Engl C, Jovanovic G, Huvet M, Toni T, Sheng X, Stumpf MPH, Buck M. 2010. Managing membrane stress: the phage shock protein (Psp) response, from molecular mechanisms to physiology. FEMS Microbiol Rev 34:797–827.

37. Ma Y, Pan C, Wang Q. 2019. Crystal structure of bacterial cyclopropane-fatty-acyl-phospholipid synthase with phospholipid. J Biochem 166:139–147.

38. Kurz M, Burch AY, Seip B, Lindow SE, Gross H. 2010. Genome-driven investigation of compatible solute biosynthesis pathways of Pseudomonas syringae pv. syringae and their contribution to water stress tolerance. Appl Environ Microbiol 76:5452–5462.

39. Stringfellow WT, Domen JK, Camarillo MK, Sandelin WL, Borglin S. 2014. Physical, chemical, and biological characteristics of compounds used in hydraulic fracturing. J Hazard Mater 275:37–54.

40. Pérez MV, Guerrero LD, Orellana E, Figuerola EL, Erijman L. 2019. Time Series Genome-Centric Analysis Unveils Bacterial Response to Operational Disturbance in Activated Sludge. mSystems 4.

41. Figueroa-Gonzalez PA, Bornemann TLV, Adam PS, Plewka J, Révész F, von Hagen CA, Táncsics A, Probst AJ. 2020. Saccharibacteria as Organic Carbon Sinks in Hydrocarbon-Fueled Communities. Front Microbiol 11:587782.

42. Ehara A, Suzuki H, Kanesaki Y, Yoshikawa H, Amachi S. 2014. Draft genome sequence of strain q-1, an iodide-oxidizing alphaproteobacterium isolated from natural gas brine water. Genome Announc 2.

43. Wu X, Spencer S, Gushgari-Doyle S, Yee MO, Voriskova J, Li Y, Alm EJ, Chakraborty R. 2020. Culturing of “unculturable” subsurface microbes: natural organic carbon source fuels the growth of diverse and distinct bacteria from groundwater. Front Microbiol 11:610001.

44. Galkiewicz JP, Kellogg CA. 2008. Cross-kingdom amplification using bacteria-specific primers: complications for studies of coral microbial ecology. Appl Environ Microbiol 74:7828–7831.

45. Quast C, Pruesse E, Yilmaz P, Gerken J, Schweer T, Yarza P, Peplies J, Glöckner FO. 2013. The SILVA ribosomal RNA gene database project: improved data processing and web-based tools. Nucleic Acids Res 41:D590–6.

46. Yilmaz P, Parfrey LW, Yarza P, Gerken J, Pruesse E, Quast C, Schweer T, Peplies J, Ludwig W, Glöckner FO. 2014. The SILVA and “All-species Living Tree Project (LTP)” taxonomic frameworks. Nucleic Acids Res 42:D643–8.

47. Andrews S. 2010. FastQC: a quality control tool for high throughput sequence data.

48. Bolger AM, Lohse M, Usadel B. 2014. Trimmomatic: a flexible trimmer for Illumina sequence data. Bioinformatics 30:2114–2120.

49. Bankevich A, Nurk S, Antipov D, Gurevich AA, Dvorkin M, Kulikov AS, Lesin VM, Nikolenko SI, Pham S, Prjibelski AD, Pyshkin AV, Sirotkin AV, Vyahhi N, Tesler G, Alekseyev MA, Pevzner PA. 2012. SPAdes: a new genome assembly algorithm and its applications to single-cell sequencing. J Comput Biol 19:455–477.

50. Arkin AP, Cottingham RW, Henry CS, Harris NL, Stevens RL, Maslov S, Dehal P, Ware D, Perez F, Canon S, Sneddon MW, Henderson ML, Riehl WJ, Murphy-Olson D, Chan SY, Kamimura RT, Kumari S, Drake MM, Brettin TS, Glass EM, Chivian D, Gunter D, Weston DJ, Allen BH, Baumohl J, Best AA, Bowen B, Brenner SE, Bun CC, Chandonia JM, Chia JM, Colasanti R, Conrad N, Davis JJ, Davison BH, DeJongh M, Devoid S, Dietrich E, Dubchak I, Edirisinghe JN, Fang G, Faria JP, Frybarger PM, Gerlach W, Gerstein M, Greiner A, Gurtowski J, Haun HL, He F, Jain R, Joachimiak MP, Keegan KP, Kondo S, Kumar V, Land ML, Meyer F, Mills M, Novichkov PS, Oh T, Olsen GJ, Olson R, Parrello B, Pasternak S, Pearson E, Poon SS, Price GA, Ramakrishnan S, Ranjan P, Ronald PC, Schatz MC, Seaver SMD, Shukla M, Sutormin RA, Syed MH, Thomason J, Tintle NL, Wang D, Xia F, Yoo H, Yoo S, Yu D. 2018. KBase: the United States Department of Energy Systems Biology Knowledgebase. Nat Biotechnol 36:566–569.

51. Gurevich A, Saveliev V, Vyahhi N, Tesler G. 2013. QUAST: quality assessment tool for genome assemblies. Bioinformatics 29:1072–1075.

52. Parks DH, Imelfort M, Skennerton CT, Hugenholtz P, Tyson GW. 2015. CheckM: assessing the quality of microbial genomes recovered from isolates, single cells, and metagenomes. Genome Res 25:1043–1055.

53. Xia M, Fu D, Chakraborty R, Singh RP, Terry N. 2019. Enhanced crude oil depletion by constructed bacterial consortium comprising bioemulsifier producer and petroleum hydrocarbon degraders. Bioresour Technol 282:456–463.

54. Pérez-Miranda S, Cabirol N, George-Téllez R, Zamudio-Rivera LS, Fernández FJ. 2007. O-CAS, a fast and universal method for siderophore detection. J Microbiol Methods 70:127–131.

55. Caporaso JG, Lauber CL, Walters WA, Berg-Lyons D, Lozupone CA, Turnbaugh PJ, Fierer N, Knight R. 2011. Global patterns of 16S rRNA diversity at a depth of millions of sequences per sample. Proc Natl Acad Sci USA 108 Suppl 1:4516–4522.

56. Bolyen E, Rideout JR, Dillon MR, Bokulich NA, Abnet CC, Al-Ghalith GA, Alexander H, Alm EJ, Arumugam M, Asnicar F, Bai Y, Bisanz JE, Bittinger K, Brejnrod A, Brislawn CJ, Brown CT, Callahan BJ, Caraballo-Rodríguez AM, Chase J, Cope EK, Da Silva R, Diener C, Dorrestein PC, Douglas GM, Durall DM, Duvallet C, Edwardson CF, Ernst M, Estaki M, Fouquier J, Gauglitz JM, Gibbons SM, Gibson DL, Gonzalez A, Gorlick K, Guo J, Hillmann B, Holmes S, Holste H, Huttenhower C, Huttley GA, Janssen S, Jarmusch AK, Jiang L, Kaehler BD, Kang KB, Keefe CR, Keim P, Kelley ST, Knights D, Koester I, Kosciolek T, Kreps J, Langille MGI, Lee J, Ley R, Liu Y-X, Loftfield E, Lozupone C, Maher M, Marotz C, Martin BD, McDonald D, McIver LJ, Melnik AV, Metcalf JL, Morgan SC, Morton JT, Naimey AT, Navas-Molina JA, Nothias LF, Orchanian SB, Pearson T, Peoples SL, Petras D, Preuss ML, Pruesse E, Rasmussen LB, Rivers A, Robeson MS, Rosenthal P, Segata N, Shaffer M, Shiffer A, Sinha R, Song SJ, Spear JR, Swafford AD, Thompson LR, Torres PJ, Trinh P, Tripathi A, Turnbaugh PJ, Ul-Hasan S, van der Hooft JJJ, Vargas F, Vázquez-Baeza Y, Vogtmann E, Caporaso JG. 2019. Reproducible, interactive, scalable and extensible microbiome data science using QIIME 2. Nat Biotechnol 37:852–857.

57. Bokulich NA, Kaehler BD, Rideout JR, Dillon M, Bolyen E, Knight R, Huttley GA, Gregory Caporaso J. 2018. Optimizing taxonomic classification of marker-gene amplicon sequences with QIIME 2’s q2-feature-classifier plugin. Microbiome 6:90.

58. Camacho C, Coulouris G, Avagyan V, Ma N, Papadopoulos J, Bealer K, Madden TL. 2009. BLAST+: architecture and applications. BMC Bioinformatics 10:421.

59. Chen I-MA, Chu K, Palaniappan K, Ratner A, Huang J, Huntemann M, Hajek P, Ritter S, Varghese N, Seshadri R, Roux S, Woyke T, Eloe-Fadrosh EA, Ivanova NN, Kyrpides NC. 2021. The IMG/M data management and analysis system v.6.0: new tools and advanced capabilities. Nucleic Acids Res 49:D751–D763.

60. Nurk S, Meleshko D, Korobeynikov A, Pevzner PA. 2017. metaSPAdes: a new versatile metagenomic assembler. Genome Res 27:824–834.

61. Kang D, Li F, Kirton ES, Thomas A, Egan RS, An H, Wang Z. 2019. MetaBAT 2: an adaptive binning algorithm for robust and efficient genome reconstruction from metagenome assemblies https://doi.org/10.7287/peerj.preprints.27522v1.

62. Olm MR, Brown CT, Brooks B, Banfield JF. 2017. dRep: a tool for fast and accurate genomic comparisons that enables improved genome recovery from metagenomes through de-replication. ISME J 11:2864–2868.

63. Chaumeil P-A, Mussig AJ, Hugenholtz P, Parks DH. 2019. GTDB-Tk: a toolkit to classify genomes with the Genome Taxonomy Database. Bioinformatics https://doi.org/10.1093/bioinformatics/btz848.

64. Langmead B, Salzberg SL. 2012. Fast gapped-read alignment with Bowtie 2. Nat Methods 9:357–359.

65. Edgar RC. 2004. MUSCLE: a multiple sequence alignment method with reduced time and space complexity. BMC Bioinformatics 5:113.

66. Tamura K, Nei M, Kumar S. 2004. Prospects for inferring very large phylogenies by using the neighbor-joining method. Proc Natl Acad Sci USA 101:11030–11035.

67. O’Leary NA, Wright MW, Brister JR, Ciufo S, Haddad D, McVeigh R, Rajput B, Robbertse B, Smith-White B, Ako-Adjei D, Astashyn A, Badretdin A, Bao Y, Blinkova O, Brover V, Chetvernin V, Choi J, Cox E, Ermolaeva O, Farrell CM, Goldfarb T, Gupta T, Haft D, Hatcher E, Hlavina W, Joardar VS, Kodali VK, Li W, Maglott D, Masterson P, McGarvey KM, Murphy MR, O’Neill K, Pujar S, Rangwala SH, Rausch D, Riddick LD, Schoch C, Shkeda A, Storz SS, Sun H, Thibaud-Nissen F, Tolstoy I, Tully RE, Vatsan AR, Wallin C, Webb D, Wu W, Landrum MJ, Kimchi A, Tatusova T, DiCuccio M, Kitts P, Murphy TD, Pruitt KD. 2016. Reference sequence (RefSeq) database at NCBI: current status, taxonomic expansion, and functional annotation. Nucleic Acids Res 44:D733–45.

68. Cantalapiedra CP, Hernández-Plaza A, Letunic I, Bork P, Huerta-Cepas J. 2021. eggNOG-mapper v2: Functional Annotation, Orthology Assignments, and Domain Prediction at the Metagenomic Scale. Mol Biol Evol 38:5825–5829.

69. Eren AM, Kiefl E, Shaiber A, Veseli I, Miller SE, Schechter MS, Fink I, Pan JN, Yousef M, Fogarty EC, Trigodet F, Watson AR, Esen ÖC, Moore RM, Clayssen Q, Lee MD, Kivenson V, Graham ED, Merrill BD, Karkman A, Blankenberg D, Eppley JM, Sjödin A, Scott JJ, Vázquez-Campos X, McKay LJ, McDaniel EA, Stevens SLR, Anderson RE, Fuessel J, Fernandez-Guerra A, Maignien L, Delmont TO, Willis AD. 2021. Community-led, integrated, reproducible multi-omics with anvi’o. Nat Microbiol 6:3–6.

70. Jain C, Rodriguez-R LM, Phillippy AM, Konstantinidis KT, Aluru S. 2018. High throughput ANI analysis of 90K prokaryotic genomes reveals clear species boundaries. Nat Commun 9:5114.

71. Pal C, Bengtsson-Palme J, Rensing C, Kristiansson E, Larsson DGJ. 2014. BacMet: antibacterial biocide and metal resistance genes database. Nucleic Acids Res 42:D737–43.

72. Oliveira JS, Araújo W, Lopes Sales AI, Brito Guerra A de, Silva Araújo SC da, de Vasconcelos Atr, Agnez-Lima LF, Freitas AT. 2015. BioSurfDB: knowledge and algorithms to support biosurfactants and biodegradation studies. Database (Oxford) 2015.

73. Blin K, Shaw S, Kloosterman AM, Charlop-Powers Z, van Wezel GP, Medema MH, Weber T. 2021. antiSMASH 6.0: improving cluster detection and comparison capabilities. Nucleic Acids Res 49:W29–W35.

