## Supplementary material for "*Iodidimonas*, a bacterium unable to degrade hydrocarbons, thrives in a bioreactor treating oil and gas produced water"

**Members of *Iodidimonas* dominate a bioreactor treating produced water despite the inability to degrade hydrocarbons**

**Material and methods**

*Artificial Pore water Media (APM) recipe:*

***Chemical***	***Amount/1000mL***

Sodium Chloride, NaCl	20g

Potassium Chloride, KCl	0.67g

DL minerals	10 mL

DL Vitamins	10 mL

RST minerals	20 mL

phosphate buffer (0.1M, pH=7)	100 mL

All the above salts and solutions were dissolved and made up to 850 mL with MilliQ water.

***Salt Stock***

Magnesium Chloride, MgCl_2_ x 6H_2_O	21.2g/100mL

Calcium Chloride, CaCl_2_ x 2 H_2_O	3.04g/100mL

Salt stock was autoclaved separately and 50 mL of it was added to the remaining media such that the total volume was 900 mL (10% volume is dedicated to inoculum addition)

***C-substrate utilization test***

**Table S1** List of single carbon substrates (10 mM) on which growth of the strains MBR-14, MBR-22, and MBR-55 were tested

| Arabinose | Glucose | L-Citrulline | Rhamnose | Ectoine |
| --- | --- | --- | --- | --- |
| Citric acid | b-lactose | L-Glutamine | Ribose | Glucose-6-phosphate |
| Cytosine | Maltose | L-Histidine | Sodium acetate | Sodium butyrate |
| D-(+)-Cellobiose | Myo-Inositol | L-Isoleucine | Sodium propionate | Sodium formate |
| D-(+)-Glucosamine Hydrochloride | DL-Proline | L-Leucine | Sodium pyruvate | N-Acetyl Glucosamine (NAG) |
| D-(+)-Mannose | D-Serine | L-Methionine | Sodium succinate (dibasic hexahydrate) | Glycerol |
| D-Mannitol | D-Alanine | L-Phenylalanine | Sucrose | D-glucuronic acid sodium salt |
| D-Sorbitol | L-Asparagine | L-Tryptophan | Thymidine | 4-hydroxybenzoate |
| DL-Malic acid (disodium salt) | L-Glutamic acid (monopotassium salt) | L-Valine | Xylitol | Sodium lactate |
| D(-)-fructose | L-Aspartic Acid | Inosine | Xylose | Ethanol |

***Hydrocarbon degradation potential***

The bacterial isolates were grown to mid-log phase in marine broth spiked with benzene, toluene, ethylbenzene, *m-, o-, p-*xylenes and naphthalene (BTEXN) and hexadecane (HD) to induce enzymes, washed thrice and resuspended in APM to be used as inoculum. To test the ability to degrade hydrocarbons, the three strains were inoculated in triplicates into 50 mL of APM in 250 mL glass serum bottles and spiked with 20 mg/L of each BTEXN or 150 mg/L of HD. Serum bottles were stoppered with butyl rubber stoppers and crimped with aluminum caps and incubated at 30 °C with no shaking. 5 mL of sample were withdrawn for hydrocarbon measurement (4 mL for Liquid-liquid-extraction with 2 mL diethyl ether for GC-MS analysis) and cell counting (1 mL for flow cytometer and OD) after introducing equal volume of filtered air. Duplicate abiotic controls amended with 1 g/L sodium azide and triplicate controls with no inoculum (uninoculated control) were also analyzed to account for abiotic removal of hydrocarbons. Additionally, positive controls containing 10 mM of sucrose instead of hydrocarbons were also incubated with each strain. Bottles amended with BTEXN were sampled on 0, 4, 7, and 30 days whereas ones with HD were sampled on 0,7, and 30 days after inoculation.

Samples (4 mL) of culture broth for analysis of hydrocarbons were collected at regular time intervals and extracted with diethyl ether (2 ml) by vigorous shaking for 2 mins. The ether layer was then collected and dried over anhydrous sodium sulfate to remove any traces of water. This ether extract was then transferred into 2mL amber GC vials and capped prior to injection via autosampler into gas chromatograph- mass spectrometer (GC-MS) for analysis. Analysis of ether extracts was performed using a gas chromatograph (Agilent GC6890N) equipped with a mass spectrometry detector (Agilent MS 5975C) with electron ionization and column used was HP-5MS (Agilent, USA). For BTEXN analysis, the inlet port was set at 250 °C with a 1:10 split ratio, and helium gas (carrier) was flowing through the column at a constant rate of 1 mL/min. The oven temperature was initially set at 35 °C for 3 min and subsequently ramped at a rate of 10 °C/min to a final temperature of 170 °C and held at 170 °C for 3 mins. For HD analysis, the inlet port was set at 250°C in a splitless mode, and helium was set at a flow rate of 1.2 mL/min. Column temperature ramped at a rate of 10°C/min to a final temperature of 300°C after an initial 1-min hold at 60°C. The hold time at the final temperature was 3 min. In all cases, mass spectra data were compared with the standard National Institute of Standards and Technology (NIST) library for compound identification using the MassHunter software.

Flow cytometric counting was performed using Attune NXT Flowcytometer (ThermoFisher Scientific, USA) equipped with an autosampler. Samples were diluted in phosphate buffer saline, stained with 1X SYBR Green I (Invitrogen, ThermoFisher) in a 96-well plate and incubated in dark for 15 mins at room temperature. 40 μL (total volume of 100 μL) of stained sample was acquired at the flow rate of 0.5 mL/min. The instrument threshold was set to 1300 on BL1 (green fluorescence) channel and 1800 on BL3 (red fluorescence) channel. Gates for bacterial counts were created on BL3 vs BLI channel such that background noise was eliminated using uninoculated control.

***Electron microscopy***

Cultures grown to log-phase on Marine broth at 30 °C for 3 days were used for electron microscopy imaging. In preparing a specimen for imaging, 4 µl of culture was applied to a glow-discharged EM grid (Formvar-carbon, 200 mesh copper, Electron Microscopy Sciences). Following five minutes the sample was partially blotted (from the side with filter paper) to leave about 1 µl and the grid was placed sample-side down on a 75 µl water drop situated on laboratory film (Parafilm M, Pechiney Plastic Packaging). The grid was transferred to a fresh drop of water after 10 seconds. This step was repeated once more after which the grid was removed from the final water drop, oriented sample-side up and partially blotted to about 1 µl to which 2 µl of 1% uranyl acetate was added. The remaining liquid was completely blotted after 10 seconds. EM grids were examined in a JEOL JEM-1200x electron microscope operating at 80 kV. A 2k x 2k pixel CCD camera (UltraScan, Gatan) controlled by the Digital Micrograph software package (Gatan) was used to record images over a magnification range of 5,000X to 30,000X.

***Emulsification Activity Measurement***

The emulsification activities of the three isolates were measured as previously described in Xia et al, 2019. Briefly, the isolates were grown in liquid APM medium amended with 10 mM glucose. Exponentially growing cultures were passed through sterile 0.2 mm syringe filters and 4 mL of cell free supernatant was combined with 4 mL of either crude oil (sourced from “North Sea Oil Field”), gasoline (Chevron), or diesel (Conoco Phillips) in round bottom graduated culture tubes. The tubes were then vortexed at maximum speed for 2 minutes to ensure thorough mixing and incubated at room temperature for 24 hours. An emulsification index, E24 (%), was calculated using the formula:

$$E24 (\%) = (\frac{HE}{HT})*100$$

where HE refers to the height of the emulsification layer (mm), and HT refers to the total height of the liquid column (mm). Control incubations were performed by mixing sterile APM media amended with 10 mM glucose with either crude oil, gasoline, or diesel at equal proportions and vortexing as described above. All incubations were performed in triplicate.

***Siderophore Production Assay(o-CAS assay)***

The isolates were grown on solid Marine Agar 2216 (MA), R2A amended with 3% NaCl, and APM amended with 10 mM glucose at 30 °C. Once visible colonies formed, the plates were overlaid with a medium designed to detect the production of siderophores using a modified protocol originally described in Pérez-Miranda et al, 2007. The overlay medium was prepared using a modified version of the protocol described in Louden et al, 2011. For 1 liter of overlay medium, 10 mL of 0.270 g/L FeCl_3_·6H_2_O, 40 mL of 1.823 g/L hexadecyltrimethylammonium bromide (HDTMA), 40.32 g/L PIPES, and 9 g/L Agar were combined in 900 mL MQ H_2_O. The medium was adjusted to pH 7 using 1M HCl and then autoclaved. After cooling down to 70 °C, 50 mL of filter sterilized 1.210 g/L Chromeazurol S was added, turning the medium blue. Approximately 25 mL of this overlay medium was poured over each plate and allowed to solidify. Plates were incubated at 30 °C for three days and were then examined for a color change from blue to orange in the regions surrounding the colonies, indicating the production of siderophores. Negative control assays were performed in parallel on plates that were not inoculated with microbes. Positive control assays were performed using a strain of *Paraburkholderia* known to produce siderophores streaked onto R2A medium and incubated at 30 °C. All assays were performed in triplicate.

**RESULTS**

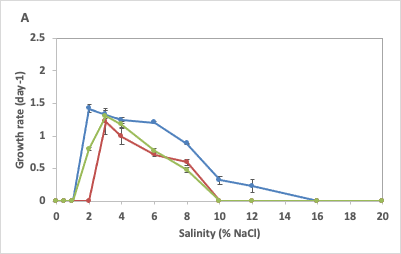

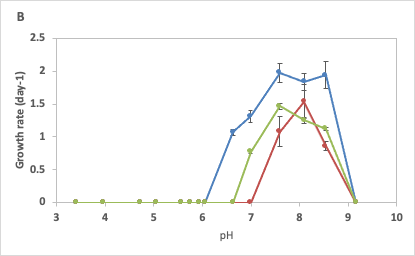

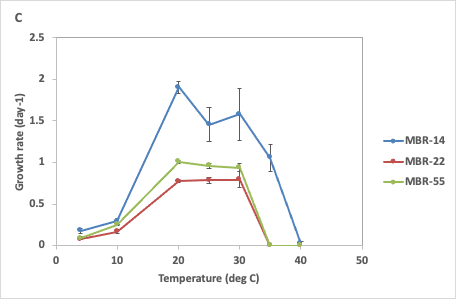

**Fig S1** (A) Salinity, (B) pH and (C) temperature growth curves for three *Iodidimonas* strains.

**Table S2** Carbon substrate utilization pattern for *Iodidimonas spp.* MBR-14, MBR-22, and MBR-55

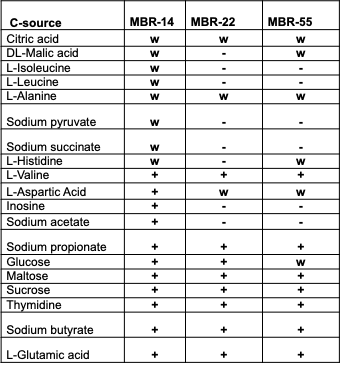

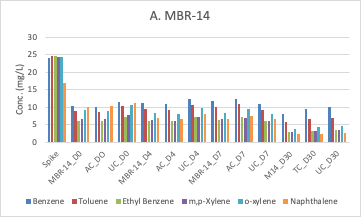

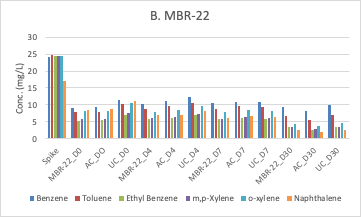

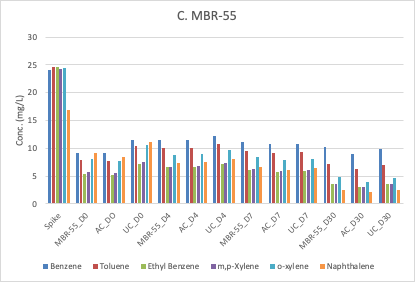

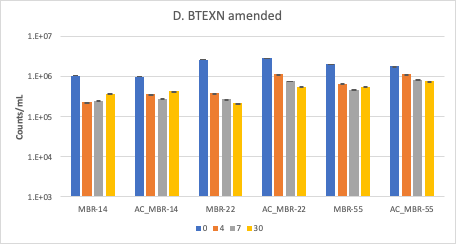

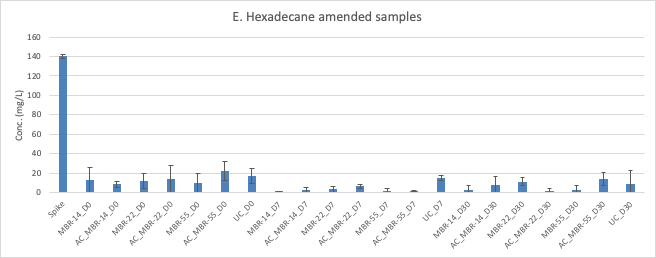

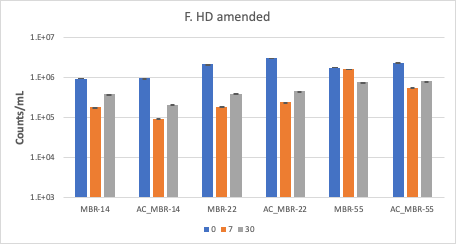

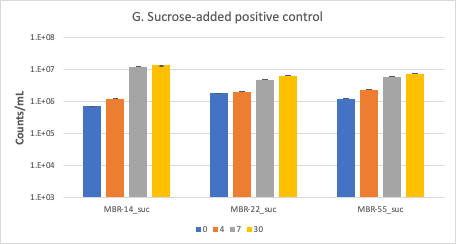

**Fig S2** Graphs showing results for BTEXN and hexadecane degradation potential tests for the three Iodidimonas strains. Graphs A, B, C show concentrations of BTEXN in liquid phase of vials inoculated with three strains (MBR-14, MBR-22, and MBR-55), abiotic control (AC) and uninoculated (UC) over 0, 4, 7 and 30 days of incubation (labels D0, D4, D7 and D30 respectively). Graph D shows flowcytometric counts of BTEXN amended vials at different days. Graph E shows concentration of hexadecane (HD) in liquid phase of vials inoculated with the three strains (MBR-14, MBR-22 and MBR-55), abiotic control (AC) and uninoculated (UC) over 0, 7 and 30 days of incubation (labels D0, D7 and D30 respectively). HD is barely soluble in water and hence subsampling lead to poor recovery of the HD in the liquid phase. D oisGraph F shows flowcytometric counts of HD-amended vials at different days. Graph G shows flowcytometric counts of positive control amended with sucrose at different days. Based on the hydrocarbon degradation and respective flowcytometric counts, the three MBR strains are not capable of degrading added hydrocarbons.

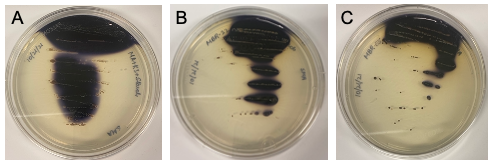

**Fig S3** Results from iodide oxidation tests performed with the three *Iodidimonas* isolates (A) MBR-14, (B) MBR-22 and (C) MBR-55 grown on MA amended with potassium iodide and starch. The formation of purple pigment around the colonies indicates conversion of iodide to molecular iodine, which in turn complexes with starch to produce the purple color.

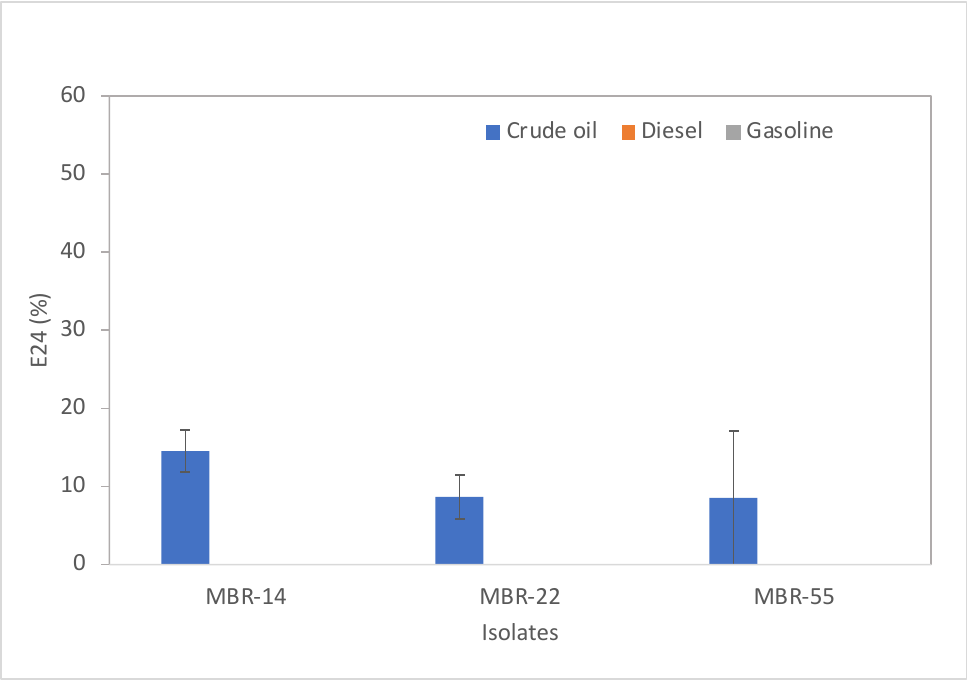

**Fig S4** Results from emulsification assays performed with the three *Iodidimonas* isolates using either crude oil, diesel, or gasoline. Error bars represent one standard deviation of experimental triplicates.

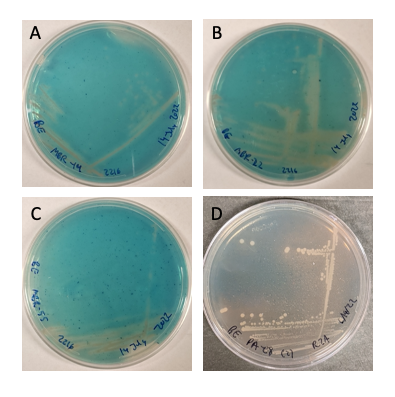

**Fig S5** Results from siderophore overlay assays with the three *Iodidimonas* isolates (A) MBR-14, (B) MBR-22, (C) MBR-55 grown on MA. The overlay assay was also performed on (D) *Burkholderia sp*. strain PA-E8, a known siderophore producing bacterium, as a positive control. A color transformation from blue to orange in areas surrounding colonies indicate the production of siderophores.

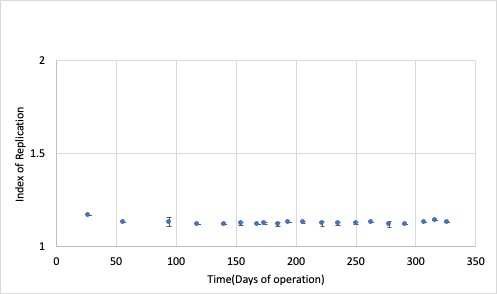

**Fig S6** Index of replication calculated from the metagenomes using reads mapped to the MAG.

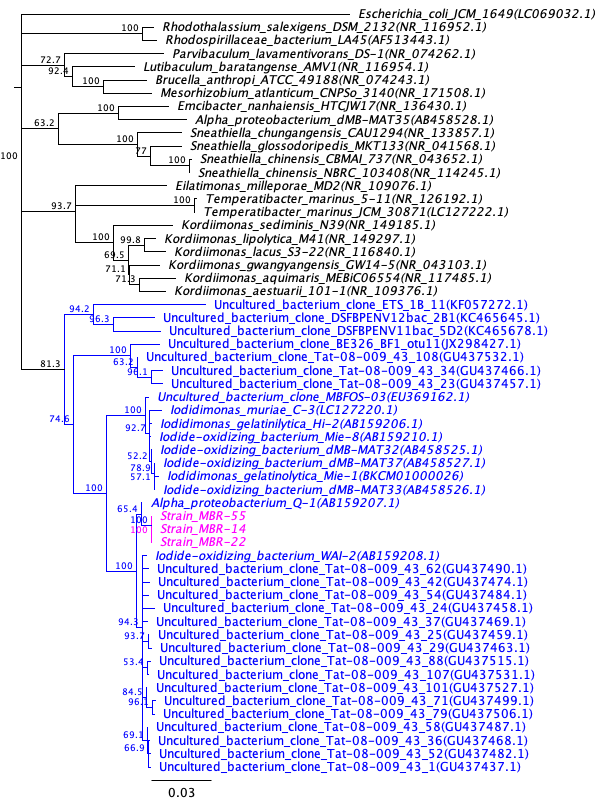

**Fig S7** Expanded phylogenetic trees based on 16S rRNA sequences (Neighbor-Joining tree); isolates or clone sequences classified as genus *Iodidimonas* in SILVA database are marked in blue and the strains from this study in pink.

**Table S3** Pairwise ANI (upper triangle) and AF (lower triangle) for the members of genus *Iodidimonas*

| **AF/ANI** | **MBR-14** | **MBR-22** | **MBR-55** | **MAG** | **Q-1** | **C-3^T^** | **Hi-2^T^** | **Mie-1** |
| --- | --- | --- | --- | --- | --- | --- | --- | --- |
| MBR-14 |  | 99.986 | 99.984 | 99.984 | 95.789 | 78.588 | 78.558 | 78.611 |
| MBR-22 | 0.985 |  | 99.997 | 99.996 | 95.784 | 78.517 | 78.559 | 78.564 |
| MBR-55 | 0.984 | 0.997 |  | 99.999 | 95.806 | 78.584 | 78.629 | 78.651 |
| MAG | 0.951 | 0.964 | 0.959 |  | 95.842 | 78.549 | 78.572 | 78.582 |
| AB Q-1 | 0.891 | 0.904 | 0.900 | 0.897 |  | 78.607 | 78.458 | 78.702 |
| C-3^T^ | 0.444 | 0.450 | 0.450 | 0.445 | 0.437 |  | 95.886 | 95.851 |
| Hi-2^T^ | 0.445 | 0.454 | 0.449 | 0.446 | 0.459 | 0.898 |  | 99.418 |
| Mie-1 | 0.465 | 0.460 | 0.447 | 0.450 | 0.467 | 0.901 | 0.938 |  |

*Complete names: *Iodidimonas sp.* MBR-14, *Iodidimonas sp.* MBR-22, *Iodidimonas sp.* MBR-55, *Iodidimonas sp.* Q-1, *Iodidimonas muriae* C-3^T^, *Iodidimonas gelatinolytica sp*. Hi-2^T^, *Iodidimonas gelatinolytica sp.* Mie-1
